# NAP: an open-source pipeline for cross-domain microbiome profiling using Nanopore sequencing-derived amplicon data

**DOI:** 10.64898/2026.05.22.727110

**Authors:** Luke B. Jones, Stefan Bagby

**Affiliations:** Department of Life Sciences, University of Bath, Bath, United Kingdom

**Keywords:** Amplicon sequencing, Microbiome, Nanopore sequencing, Ribosomal RNA, Taxonomic classification

## Abstract

**Background:** Nanopore sequencing offers a cost-effective and portable platform for microbiome analysis, but amplicon-based approaches remain limited by higher sequencing error rates and a lack of workflows tailored to mixed domain ribosomal RNA profiling. While short-read technologies dominate microbial community analysis, their portability and flexibility are constrained. There is therefore a need for robust pipelines designed specifically for cross-domain Nanopore amplicon data.

**Results:** We introduce the Nanopore sequencing-based Amplicon Pipeline (NAP; https://github.com/Luke-B-Jones/NAP), an open-source workflow optimised for flexible mixed domain primer sets such as 515Y/926R. NAP performs adaptive quality filtering, chimera removal, centroid generation, BLAST-based taxonomic classification, hierarchical consensus correction, and domain-aware post-processing, outputting decontaminated abundance tables suitable for downstream analysis. Initial validation against two complementary commercial mock communities showed that NAP achieved strong genus-level performance across both low complexity logarithmic and more compositionally complex gut mock communities. Detection was most reliable above *ca.* 1% relative abundance, and replicate outputs showed strong agreement with expected composition under Bray–Curtis, Jaccard, agreement-plot, and Bland–Altman analyses. Benchmarking of NAP’s internal filtering modes showed that the default adaptive setting provided the most robust balance of read quality, retained depth, and downstream taxonomic fidelity across heterogeneous inputs. Direct comparison against QIIME2 and Kraken2/Bracken further showed that NAP most accurately preserved expected community structure, with markedly fewer false positive assignments at genus level and substantially stronger species-level behaviour under the tested conditions. Species-level assignments were informative for some taxa, but remained less robust than genus-level outputs with the default V4–V5 amplicon.

**Conclusions:** NAP provides a robust and flexible workflow for cross-domain Nanopore amplicon profiling, with strongest performance at genus level and competitive species-level behaviour for well resolved taxa. Although analysis of field-derived data was not assessed here, NAP compatibility with portable Nanopore sequencing supports accurate mixed domain microbiome profiling under the tested conditions.

## Background

Microbiomes are increasingly studied from a cross-domain perspective, capturing interactions among bacteria, archaea, and eukaryotes across diverse environmental and host-associated contexts. While microbiome research has historically focused on bacteria, recent work highlights that inter-domain interactions are central to understanding community structure and function, and health-related outcomes. In clinical settings, bacterial–fungal interactions have physical, metabolic, and immune implications in disease (Arvanitis and Mylonakis, 2015). Inter-kingdom microbial networks are now recognized as potential drivers of complex pathologies and may offer novel diagnostic and therapeutic targets (Li *et al*., 2018; Wang *et al*., 2023). Beyond medicine, these interactions shape ecological dynamics, influence food microbiology, and affect agricultural productivity (Frey-Klett *et al*., 2011). As such, accurate and domain-inclusive profiling of microbial communities is essential for advancing microbiome science across multiple disciplines.

While shotgun and whole genome sequencing approaches offer high taxonomic resolution and low taxonomic bias, they remain costly and computationally intensive. For studies focused primarily on community composition, amplicon sequencing provides a faster and more cost-effective alternative. Traditionally, however, amplicon-based profiling has been restricted to bacteria using 16S rRNA primers, or has relied on separate 16S and 18S reactions to study bacterial and eukaryotic components independently. Because these are amplified and sequenced in isolation, their abundance outputs are not directly comparable, limiting cross-domain integration.

Recent advances in primer design have addressed this issue by enabling the amplification of bacterial, archaeal, and eukaryotic small subunit rRNA from a single reaction using universal primer sets. Illumina-based workflows offer high base accuracy and are compatible with such primers, but remain limited by short read lengths. Moreover, while mature and user-friendly pipelines exist for Illumina data, these often require programming expertise and lack tools for domain-aware normalisation or bias correction. Nanopore sequencing, in contrast, supports real-time, full length amplicon sequencing with much lower capital outlay and greater portability (Lao *et al*., 2021; Charalampous *et al*., 2019). These features make it an appealing choice for field-based or resource-limited settings. However, existing Nanopore amplicon tools are largely limited to bacterial 16S pipelines (e.g., EPI2ME) or require researchers to build bespoke workflows from scratch, rendering them inaccessible to most potential users, particularly those aiming to study mixed domain microbial communities.

We present here the Nanopore sequencing-based Amplicon Pipeline (NAP), a lightweight, open-source pipeline purpose built to transform Nanopore amplicon sequencing data into high quality, taxonomically normalised microbiome profiles. To our knowledge, no other end-to-end pipeline currently exists for the analysis of cross-domain ribosomal RNA amplicon data generated using Nanopore sequencing. NAP addresses this gap by integrating established sequence processing tools with custom algorithmic components into a single workflow that can accommodate cross-domain rRNA primer sets (such as the default set, 515Y/926R), supporting simultaneous bacterial, archaeal, and eukaryotic profiling from a single amplicon pool.

## Implementation

NAP is implemented as an end-to-end command line workflow in Bash and Python for Unix/Linux-based execution, with Docker support to simplify installation and improve reproducibility across systems. Established third party tools are used for core sequence processing steps, whilst custom Bash wrappers and Python modules handle workflow orchestration, taxonomy parsing, consensus calling, abundance processing, decontamination, and output summarisation (Figure 1). This structure was adopted to preserve a simple user interface whilst retaining modularity at code level, allowing individual components to be updated or replaced without altering the overall workflow.

**Figure 1:**
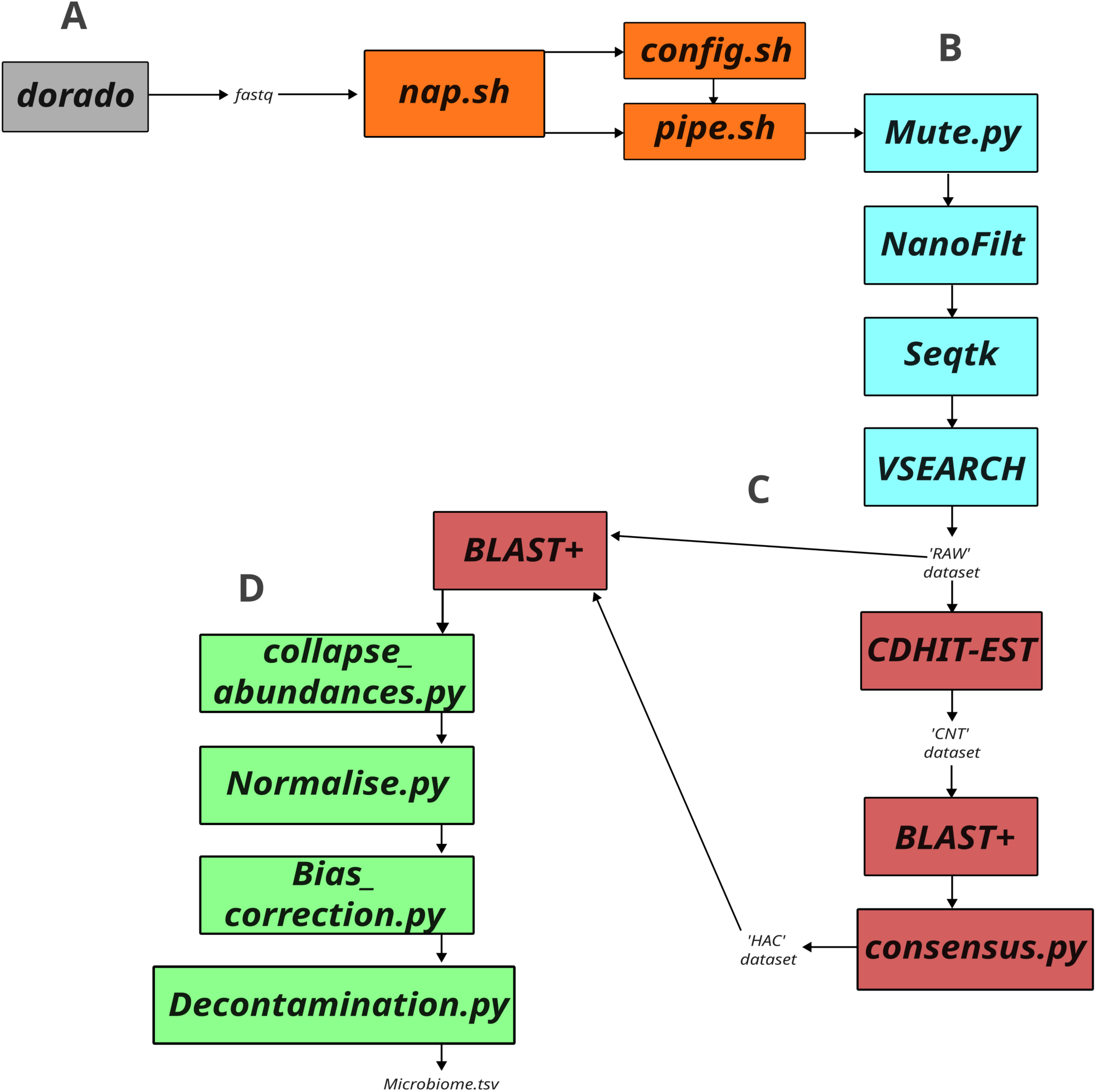
Core pipeline workflow. (A) Dorado output is the input to the pipeline, and is fed into the wrapper nap.sh which passes variables along with the config to pipe.sh, initiating the pipeline. (B) Quality control stages shown in light blue. (C) ‘RAW’ data are clustered and the output centroids (‘CNT’ dataset) are then taxonomically classified. Resulting taxonomic classification data from CNT are synthesised into a ‘higher accuracy consensus’ (HAC) dataset, against which raw data are then BLASTed, applying an abundance dimension to HAC taxonomy. (D) Final BLAST outputs are simplified to genus and species level, collapsing taxonomy, and combining abundances accordingly. This dataset is then normalised using total sum scaling (TSS) normalisation. If blanks are set up correctly (using ‘nap decon’), Decontamination.py then removes organisms based on blank content, outputting Microbiome.tsv. Tools used: NanoFilt (v2.8.0; De Coster *et al.,* 2018), SeqTK (v1.4; Li, 2026), VSEARCH (v2.28.1; Rognes *et al*., 2016), BLAST (v2.16.0; Camacho *et al*., 2009), and CD-HIT (4.8.1; Li and Godzik, 2006).

SILVA rRNA reference databases are first refined to remove ambiguous, taxonomically unresolved, off-target, and redundant entries, and in the default ‘mammalian_microbiome_inclusive’ mode, contaminant and organellar records such as plant mitochondrial rRNA are also removed. Alternative filtering modes can also be defined by the user. The curated reference is then divided into 16S and 18S subsets so that primer mapping and amplicon extraction can be performed within the expected domain-specific sequence space, accounting for the different amplicon structures generated by mixed domain primer sets. Primer sequences are mapped to each retained entry using ungapped local alignment, and the longest predicted amplicon region is extracted where possible; untrimmed entries are retained only where strict trimming would result in unnecessary loss of reference coverage. These primer-aware, domain-specific reference sets are then indexed for BLAST-based classification. This design reduces redundant high-scoring matches, limits ambiguous search space, and decreases processing time without requiring users to manually construct primer-specific databases for each assay.

NAP processes demultiplexed amplicon reads (FASTQ) through a centroid-first workflow optimised for mixed domain Nanopore microbiome profiling. Reads are first subjected to adaptive quality control using an input depth-dependent filtering scheme. Samples are assigned to preset read depth bins according to the initial number of input reads, with each bin linked, *via* the NAP configuration file, to a corresponding read level Phred threshold and base masking stringency. Under the validation conditions used here, these settings were empirically calibrated to favour the highest average read quality whilst retaining ≥100,000 reads where possible. Firstly, base qualities are tallied across the dataset, and a masking cutoff is selected such that only a defined high quality proportion of bases remains unmasked; bases below this cutoff are converted to ‘`N’` rather than causing the entire read to be discarded in downstream mean Phred filtering. Following this, the associated Phred is applied. This design allows higher depth datasets to be filtered more stringently while preserving usable sequence in lower depth samples. The exact default bin definitions used here are listed in Supplementary Table 1. Users can modify these thresholds to better suit alternative datasets. Once filtering has been completed, very deep samples are capped by default at 225,000 reads to prevent unnecessary computational overhead during downstream clustering and classification. Chimeric reads are then removed with VSEARCH, and the filtered read set (‘RAW’) is clustered with CD-HIT to generate representative centroids (‘CNT’). These centroid sequences are classified against the primer-trimmed SILVA database using BLAST+, with alignment settings tuned separately for centroid-to-reference classification and subsequent RAW-to-centroid reassignment, reflecting the distinct alignment objectives of these stages. Taxonomic consensus is then derived from the retained BLAST hits using a hierarchical frequency-weighted procedure. Briefly, the dominant taxonomic structure across the top matches is identified, and the best supported representative assignment is then selected using percent identity, bit score, and e-value. The annotated centroid set (‘HAC’) is subsequently converted into an internal high confidence database against which all RAW reads are re-aligned, preserving abundance information whilst reducing sensitivity to noisy single read classifications and limiting overall runtime.

Post-processing is performed automatically on the resulting abundance tables. Taxonomic assignments can be collapsed to genus and species level, normalised using total sum scaling (TSS), and filtered to remove low confidence taxa below the reporting threshold. For the bundled 515Y/926R configuration, NAP applies a default 16S/18S correction factor of 0.4 to account for the reduced amplification efficiency of co-amplified 18S templates relative to 16S targets. When matched blank controls are supplied, and decontamination is activated in the configuration, blank-informed decontamination is performed by temporarily denormalising and harmonising sample and blank abundance profiles across taxonomic levels, before evaluating taxa using configurable prevalence- and abundance-based rules. Taxa supported primarily by blank signal are then removed or penalised adaptively before final re-normalisation. NAP does not currently apply taxon-specific SSU copy number correction, and output abundances should therefore be interpreted as relative amplicon abundances rather than direct estimates of cellular abundance.

Global run behaviour is controlled through a main configuration file, whilst primer-specific defaults are defined in modular sub-configurations containing the expected amplicon structure, reference resources, alignment thresholds, and any domain-specific abundance adjustments. This enables users to adapt the workflow to alternative rRNA primer sets without modifying the core code. Input can be supplied either as a single explicitly pathed read file with a user-defined sample name, or as a tabular manifest for batch execution. For manifest-based execution, output locations are user-defined, allowing runs to be initiated from any working directory. Final outputs include genus- and species-level abundance tables (.tsv), pipeline logs (.txt), intermediate classification files, and quality summaries intended to support troubleshooting and user confidence, whilst also allowing users to interrogate full taxonomic paths and, if desired, make manual taxonomic consensus assessments.

## Methods

### Mock community preparation

Two commercially available mock microbiome communities were used to evaluate the pipeline: ZymoBIOMICS Microbial Community Standard II (Log Distribution), and ZymoBIOMICS Gut Microbiome Standard (both Cambridge Biosciences); henceforth, these are referred to as log mock (L1, L2, and L3) community and gut mock (M1, M2, and M3) community, respectively. These two standards were selected because they impose complementary validation challenges relevant to mixed domain Nanopore amplicon profiling. The ZymoBIOMICS Microbial Community Standard II (Log Distribution) provides a low complexity community with an extreme abundance gradient, spanning *ca*. 89.1% to 0.089% relative abundance, and is therefore well suited for assessing abundance-dependent detection, dropout, and quantitative recovery across a wide dynamic range. In contrast, the ZymoBIOMICS Gut Microbiome Standard provides a more compositionally complex community, comprising 14 taxa distributed between 14% and 0.1% relative abundance, with broader phylogenetic and microbiological diversity, including taxa expected to cover varied taxonomic separability within the V4–V5 region. Together, the two standards include bacterial, eukaryotic, and archaeal targets, encompass both Gram-positive and Gram-negative organisms, and represent taxa expected to vary in extractability, amplification behaviour, and taxonomic separability within the V4–V5 region. As such, they were chosen to test the principal analytical challenges addressed by NAP: cross-domain recovery, abundance fidelity, taxonomic assignment, and reproducibility across distinct community structures.

In addition to mock communities, environmental blanks were made by exposing sterile 0.9% saline to the laboratory environment. DNA was extracted from these samples using ZymoBIOMICS DNA/RNA miniprep kit according to the manufacturer’s recommendations (“ZymoBIOMICS DNA/RNA Miniprep Kit,” 2025). Homogenisation was performed according to a custom protocol, with six cycles of 30 seconds at 9,000 rpm using a Bertin Precellys Evolution bead beater, increasing the yield of harder to lyse organisms (Zhang *et al*., 2020). DNA was quantified using a Qubit fluorometer (Thermo Fisher Scientific). Isolates were then amplified using Phusion™ Plus PCR Master Mixes (Thermo Fisher Scientific), using an annealing temperature of 50 °C and *ca.* 100ng of template DNA *per* reaction, along with the recommended guidelines for 50 µL single reaction protocol outlined by the supplier, and custom oligonucleotide primers (Eurofins) (515Y, 5′-*GTGYCAGCMGCCGCGGTAA*, 926R, 5′-*CCGYCAATTYMTTTRAGTTT;* McNichol *et al*., 2021). All amplifications were validated using gel electrophoresis by identifying 16S and 18S bands at *ca.* 300-500 and *ca.* 700 bases, respectively. This resulted in six amplicon pools, three replicates *per* mock community. Amplicon pools were then sequenced using one MinION R10.4.1 flow cell according to the Native Barcoding Protocol (SQK-NBD114.24), and then basecalled and demultiplexed using Dorado (v0.9.0; “nanoporetech/dorado,” 2025; super accuracy model v5.0.0).

### Benchmarking

To assess the suitability of NAP’s default adaptive filtering mode, alternative fixed-Q20 and fixed-Q30 filtering regimes together with modified base masking settings were benchmarked, and downstream performance was compared using Bray–Curtis and Jaccard dissimilarity, precision, recall, and F1-score at both genus and species levels.

As no end-to-end workflow specifically designed for Nanopore-derived cross-domain 515Y/926R amplicon data was identified, benchmarking was performed against two widely used and broadly applicable taxonomic profiling frameworks: QIIME2 (qiime2-amplicon-2026.1) (Bolyen *et al*., 2019) and Kraken2 (v2.1.3) (Wood, Lu and Langmead, 2019; Lu *et al*., 2022) with Bracken (v3.0.1) (Lu *et al*., 2017). In each case, benchmarking was performed using recommended standard analytical pathways and default settings wherever possible, without parameter tuning intended to favour performance on the present dataset. SILVA 138.2 SSURef NR99-based reference resources appropriate to each framework were used for taxonomic classification. The same demultiplexed and primer/adaptor-trimmed sample set was used for all workflow comparisons.

For QIIME2, reads were imported as single-end FASTQ files *via* a manifest and processed through the recommended amplicon workflow stages for quality filtering, dereplication, chimera screening, and taxonomic classification, excluding denoising steps designed for short-read Illumina datasets. Taxonomic assignments were generated using the classify-sklearn naive Bayes approach, and resulting feature tables were collapsed to genus and species level for downstream comparison. For Kraken2/Bracken, each sample was classified using Kraken2, and genus- and species-level abundance estimates were subsequently generated from Kraken2 reports using Bracken. Resulting abundance tables were exported on a *per* sample basis for downstream comparison, and a minimum abundance threshold of 0.05% was enforced as with NAP.

### *In silico* primer bias assessment

To aid interpretation of abundance distortions potentially attributable to primer–template mismatch, the default 515Y/926R primer set was evaluated *in silico* using TestPrime against the SILVA SSU RefNR 138.2 database after redundancy pruning. Up to three mismatches were permitted, with no enforced zero mismatch zone at the 3′ end. For taxa represented in the mock communities, we summarised perfect match frequency, mean pair coverage, and the proportion of reference matches containing at least one 3′ mismatch in the forward and reverse primers. These results were used descriptively to contextualise observed under-representation and were not incorporated into taxonomic assignment or abundance correction.

### Statistical analysis

Statistical analysis was performed using a custom Python script to evaluate the accuracy, reproducibility, and community structure fidelity of the pipeline’s outputs relative to the known composition of two commercially available mock communities. Genus- and species-level taxonomic tables were generated from each sample. To enable comprehensive sensitivity profiling, taxa absent across all replicates were assigned zero counts.

Detection accuracy was quantified by calculating precision, recall, and F1-scores for each replicate at a fixed detection threshold (τ = 0.001) at both genus and species level, using the expected mock composition at the corresponding taxonomic rank as the reference set. To evaluate quantitative agreement between observed and expected taxon abundances, linear regression and Lin’s concordance correlation coefficient (*ρ*_c_) were used. This enabled both proportional and absolute agreement to be assessed. Agreement plots compared mean observed abundances across replicates against their expected values, while Bland–Altman plots quantified systematic bias and calculated limits of agreement using ±1.96 standard deviations from the mean difference. Together, these analyses assessed not only detection but also abundance accuracy and consistency. This was restricted to taxa which were expected, meaning false positives were not included in this part of the analysis.

Community-level similarity and sample structure were explored using β-diversity metrics. Pairwise dissimilarities were calculated using both Jaccard distance (based on presence/absence data) and Bray–Curtis dissimilarity (based on relative abundances). These dissimilarity matrices were visualised using principal coordinate analysis (PCoA) to identify clustering of replicates around their respective mock community centroids. A replicate was defined as a success if it was closer in β-diversity space to its own mock profile than to that of the alternate mock group. Statistical significance of replicate fidelity was assessed using one-sided binomial tests for each group, with groupwise *p*-values aggregated *via* Fisher’s method to test for global consistency. To investigate taxon-level variability, bar plots were generated showing the expected genus abundances overlaid with individual replicate values. For each genus, the coefficient of variation (CV) across replicates was annotated, with values exceeding 0.5 highlighted to indicate high within-group variability. Detection sensitivity was further examined by plotting, at both genus and species level, the proportion of replicates in which each expected taxon was detected against its expected relative abundance. These analyses included all observed taxa, including false positives.

Development-stage optimisation analyses used to select the final NAP configuration, referred to as the default here, are summarised in the Supplementary material.

## Results

### Read filtering and quality control benchmarking

To assess whether NAP’s default adaptive filtering represented an appropriate general use operating mode, we benchmarked it against fixed, non-adaptive Phred filtering (Q20 and Q30), together with modified base masking regimes (mute lowest 0%, 5% and 10% Phred bases), and evaluated downstream taxonomic fidelity using Bray–Curtis and Jaccard dissimilarity, precision, recall, and F1-score at both genus and species levels (Figure 2). The default configuration (Default mute / Dynamic Phred) remained among the strongest performing settings across both taxonomic ranks, yielding mean genus-level Bray–Curtis and Jaccard dissimilarities of 0.20 and 0.31, respectively, and mean species-level values of 0.25 and 0.40. Although it was not the single numerical best performer for every summary metric, it consistently occupied the near-optimal range across abundance-sensitive, presence/absence, and classification-accuracy measures, whilst avoiding the marked degradations observed under overly stringent or aggressively masked alternatives.

**Figure 2:**
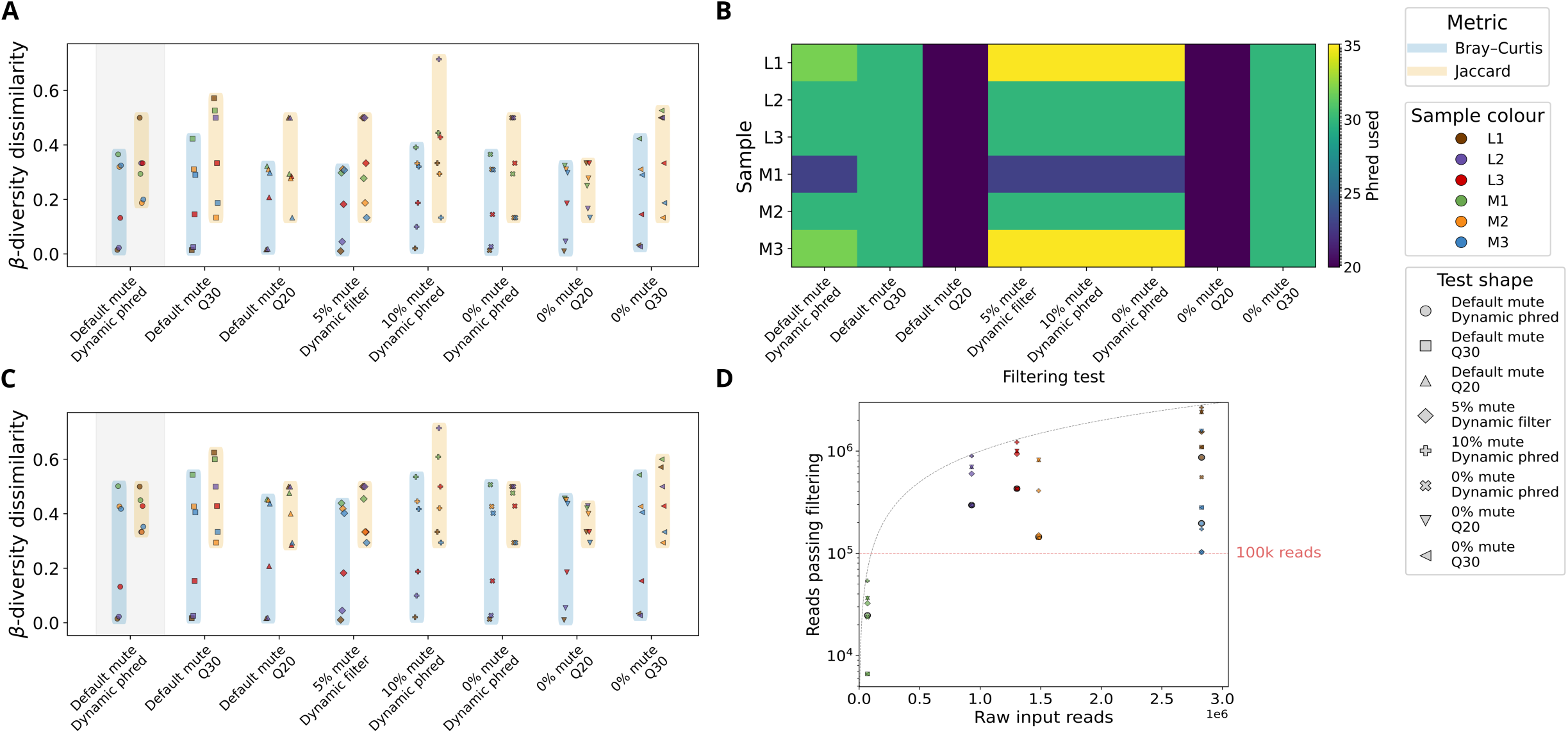
Benchmarking of NAP filtering strategies and default setting selection. (A) Genus-level β-diversity dissimilarity of benchmarked filtering modes relative to expected mock community composition. (B) Heatmap showing the Phred threshold applied to each sample under each filtering mode. (C) Species-level β-diversity dissimilarity of benchmarked filtering modes relative to expected mock community composition. (D) Reads retained after filtering plotted against raw input read count, illustrating the relationship between input depth and retained read depth under the benchmarked settings. Colours identify individual samples (L1–L3 and M1–M3), and shapes indicate the filtering mode applied.

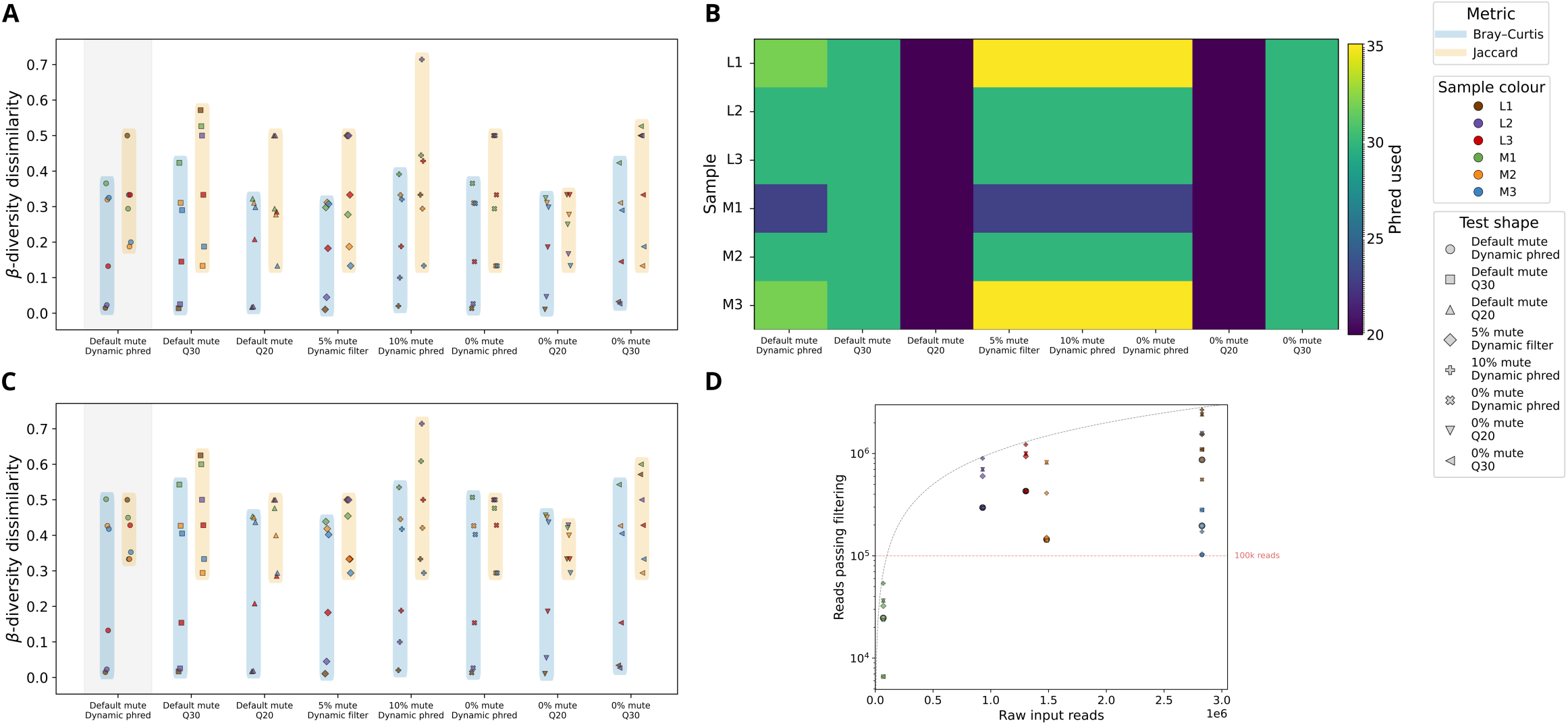

The adaptive nature of this behaviour is evident from the sample-specific Phred thresholds selected by the default workflow, which ranged from Q23 to Q35 across the six replicates (Figure 2B). High input samples (929,330–2,828,709 raw reads) were filtered at Q30–32 and retained 144,628–868,391 reads, whereas the low input replicate M1 (69,141 raw reads) was relaxed to Q23, and retained 24,699 reads, demonstrating that the default dynamic filtering strategy balanced higher read quality against retention of ≥100,000 reads wherever this was feasible (Figure 2D). In this edge case, adaptive relaxation was clearly beneficial: fixed Q30 retained only 6,603 reads for M1 and produced poorer community fidelity at both genus level (Bray–Curtis 0.37 vs. 0.42; Jaccard 0.29 vs. 0.53) and species level (Bray–Curtis 0.50 vs. 0.54; Jaccard 0.45 vs. 0.60). Across all six replicates, fixed Q30 also increased mean Jaccard dissimilarity at both genus level (0.31 vs. 0.38) and species level (0.40 vs. 0.46), indicating reduced taxonomic recovery under an overly stringent global threshold. A fixed Q20 threshold increased retention in M1 to 36,715 reads, but did not provide a uniformly superior outcome. For this sample, genus-level Bray–Curtis improved (0.32 vs. 0.37) and genus-level Jaccard remained unchanged (0.29), whilst species-level Bray–Curtis also improved (0.45 vs. 0.50) but species-level Jaccard increased slightly (0.48 vs. 0.45), indicating a mixed effect of greater retained depth and reduced read quality. The same pattern was evident across the full dataset. Although fixed Q20 increased mean retained depth more than threefold relative to the default (1,094,461 vs. 326,701 reads), it did so only by uniformly lowering all samples to Q20, and did not improve overall β-diversity fidelity relative to the adaptive default (mean genus Bray–Curtis/Jaccard 0.20/0.33 vs. 0.20/0.31; mean species Bray–Curtis/Jaccard 0.26/0.41 vs. 0.25/0.40) and therefore represented a permissive sensitivity-maximising regime rather than the most robust general use default. More aggressive masking was also clearly detrimental, with the ‘10% mute, Dynamic Phred’ mode producing the highest mean genus- and species-level Jaccard dissimilarities (0.39 and 0.48, respectively; Figure 2A,C). Classification rates were consistently high across all filtering strategies, typically exceeding 99.6% across samples, with only minor reductions observed in lower-depth datasets and under more aggressive filtering regimes. Under the default configuration, classification ranged from 98.53% to 99.88% in >Q30 samples and reached 94.68% in Q20 samples, indicating stable taxonomic assignment across variable input conditions. Taken together, these results show that although the default dynamic filtering mode is not the single numerical optimum for every benchmark endpoint, it provides the most robust balance of read quality, retained depth, and downstream taxonomic fidelity across heterogeneous inputs, and is therefore the most defensible general use setting for automated mixed domain microbiome profiling. Dataset-specific optimisation of these filtering thresholds may allow users to increase output accuracy, and is encouraged.

### Benchmarking against existing pipelines

Benchmarking NAP against QIIME2 and Kraken2/Bracken showed that all three workflows broadly recovered the expected genera, with strongest agreement in the lower complexity logarithmic mock and weaker agreement in the more compositionally complex gut mock (Figure 4A). At genus level, concordance with expected taxa was high for NAP and Kraken2 in the logarithmic mock (*ρ_c_* = 0.99 and 0.96, respectively), but lower for QIIME2 (*ρ_c_* = 0.54); the same pattern was retained in the gut mock, although with reduced concordance overall (NAP *ρ_c_* = 0.46, Kraken2 *ρ_c_* = 0.43, QIIME2 *ρ_c_* = 0.18). When restricted to expected taxa, F1-scores were therefore broadly comparable across tools, ranging from 0.76 to 0.91 with the logarithmic mock and from 0.85 to 0.94 with the gut mock. Inclusion of all detected genera, however, clearly differentiated the workflows: NAP retained high all taxa F1-scores with both the logarithmic and gut mocks (0.76 and 0.87, respectively), whereas QIIME2 declined to 0.34 and 0.41, and Kraken2 to 0.32 with both communities. This difference was driven primarily by false positive detection: across all genus-level outputs, NAP yielded 0–3 unexpected genera *per* replicate, compared with 11–32 for QIIME2 and 19–49 for Kraken2. Thus, although genus-level recovery of expected taxa was directionally similar across tools, NAP was substantially more conservative in its handling of unexpected classifications.

**Figure 4:**
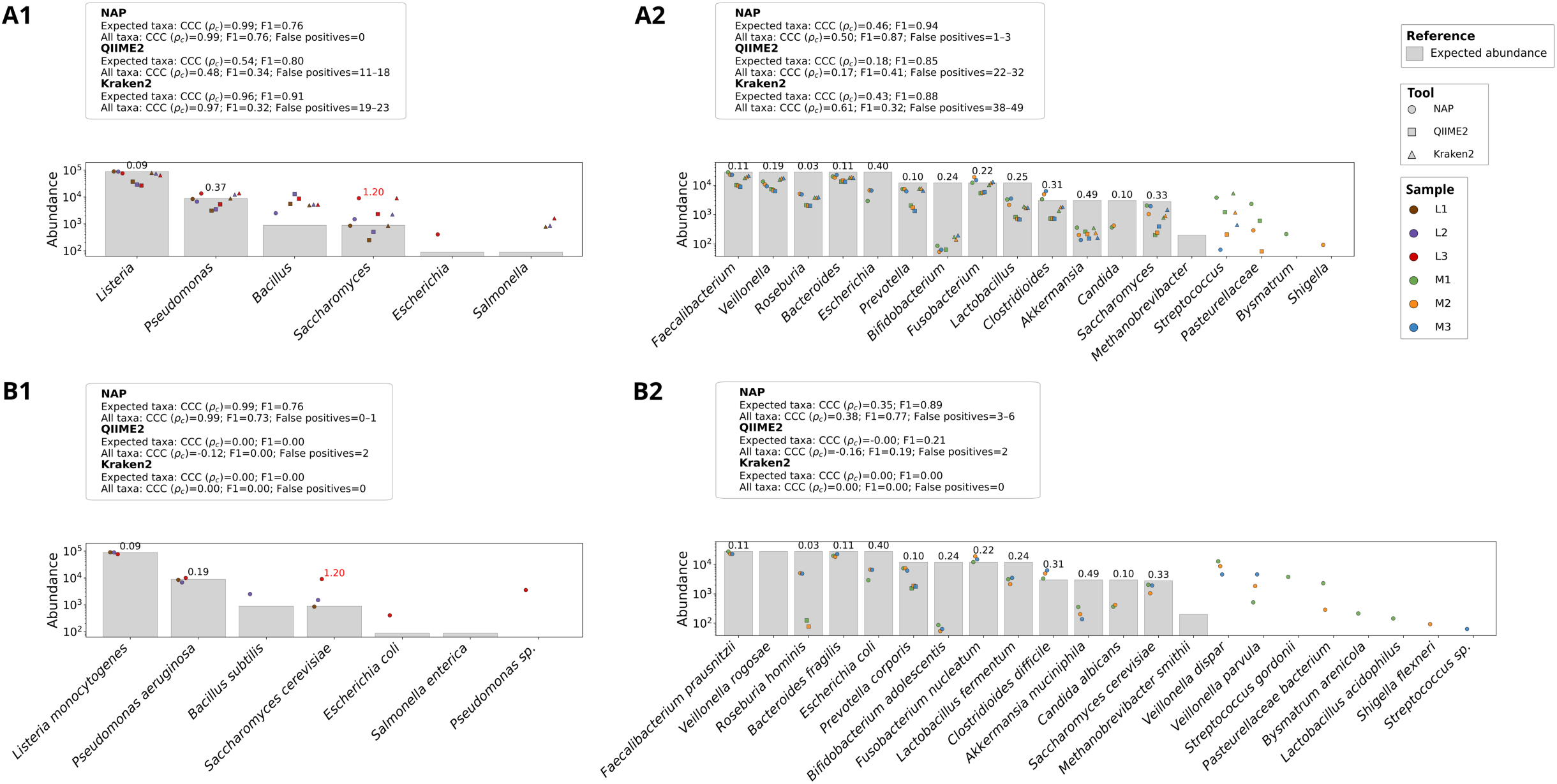
Observed taxonomic abundances across mock communities at genus and species levels. (A) Genus level results. (B) Species level results. Subpanels (1) show the gut microbiome mock (M1–M3) and subpanels (2) show the logarithmic mock (L1–L3). Grey bars represent expected abundances, while coloured points indicate observed abundances *per* replicate. Coefficient of variation for expected taxa across repeats is annotated above bars; values >0.5 are coloured red. Above each bar plot, summary metrics are provided for each workflow. The first line is calculated using expected taxa only, whereas the second includes all detected taxa and therefore incorporates contaminants and false positives. Lin’s concordance correlation coefficient (*ρ_c_*) and F1-score are shown in both cases, and the second line additionally reports the range of false positive taxa identified *per* replicate.

This contrast became much more pronounced at species level. For the logarithmic mock, NAP retained near perfect concordance with the expected species composition (*ρ_c_* = 0.99 for both expected only and all taxa analyses), whereas QIIME2 fell to 0.00 for expected taxa and –0.12 when all taxa were included, and Kraken2 returned 0.00 in both analyses. The same trend was observed in the gut mock, where NAP remained quantitatively informative (*ρ_c_* = 0.35 for expected taxa; 0.38 for all taxa), whilst QIIME2 again declined to 0.00 and –0.16, and Kraken2 remained at 0.00. F1-scores showed the same pattern: NAP degraded least from genus to species level, retaining F1 values of 0.76 and 0.73 with the logarithmic mock, and 0.89 and 0.77 with the gut mock, for expected only and all taxa analyses, respectively. In contrast, QIIME2 species-level performance collapsed to 0.00 with the logarithmic mock and remained low with the gut mock (0.21 expected taxa; 0.19 all taxa), whilst Kraken2 yielded 0.00 throughout. Inspection of the species-level outputs indicated that QIIME2 produced only sparse species assignments, often to unexpected species within otherwise expected genera, whereas Kraken2 failed to recover meaningful species-level profiles under the tested conditions. Although NAP also produced a small number of unexpected species-level assignments, these remained limited relative to the comparator workflows and were largely attributable to artefactual splitting of expected taxa or low level contamination, as discussed below.

Global β-diversity analyses supported the same interpretation (Figure 5). At genus level, NAP outputs clustered closest to the expected mock centroids under both Bray–Curtis and Jaccard dissimilarity (logarithmic mock: Bray-Curtis 0.01-0.13, Jaccard 0.33-0.50; gut mock: Bray-Curtis 0.32-0.39, Jaccard 0.19-0.29) with Kraken2 representing the nearest alternative (logarithmic mock: Bray-Curtis 0.02-0.23, Jaccard 0.80-0.83; gut mock: Bray-Curtis 0.39, Jaccard 0.80-0.84) and QIIME2 consistently the most displaced (Bray-Curtis 0.59-0.67, Jaccard 0.78-0.89; across both communities). At species level, this relationship became more extreme: QIIME2 and Kraken2 approached near-universal dissimilarity from the expected profiles (Jaccard 0.88–1.00; Bray–Curtis 0.94–1.00), whereas NAP remained substantially closer to the expected community structure (logarithmic mock: Bray–Curtis 0.01–0.13, Jaccard 0.33–0.50; gut mock: Bray–Curtis 0.42–0.50, Jaccard 0.33–0.45). Taken together, these benchmarks indicate that, under the tested conditions, NAP most accurately preserved both taxonomic composition and abundance structure, with the clearest advantages arising from reduced false positive detection at genus level and substantially improved species-level behaviour relative to the more general purpose comparator pipelines.

**Figure 5:**
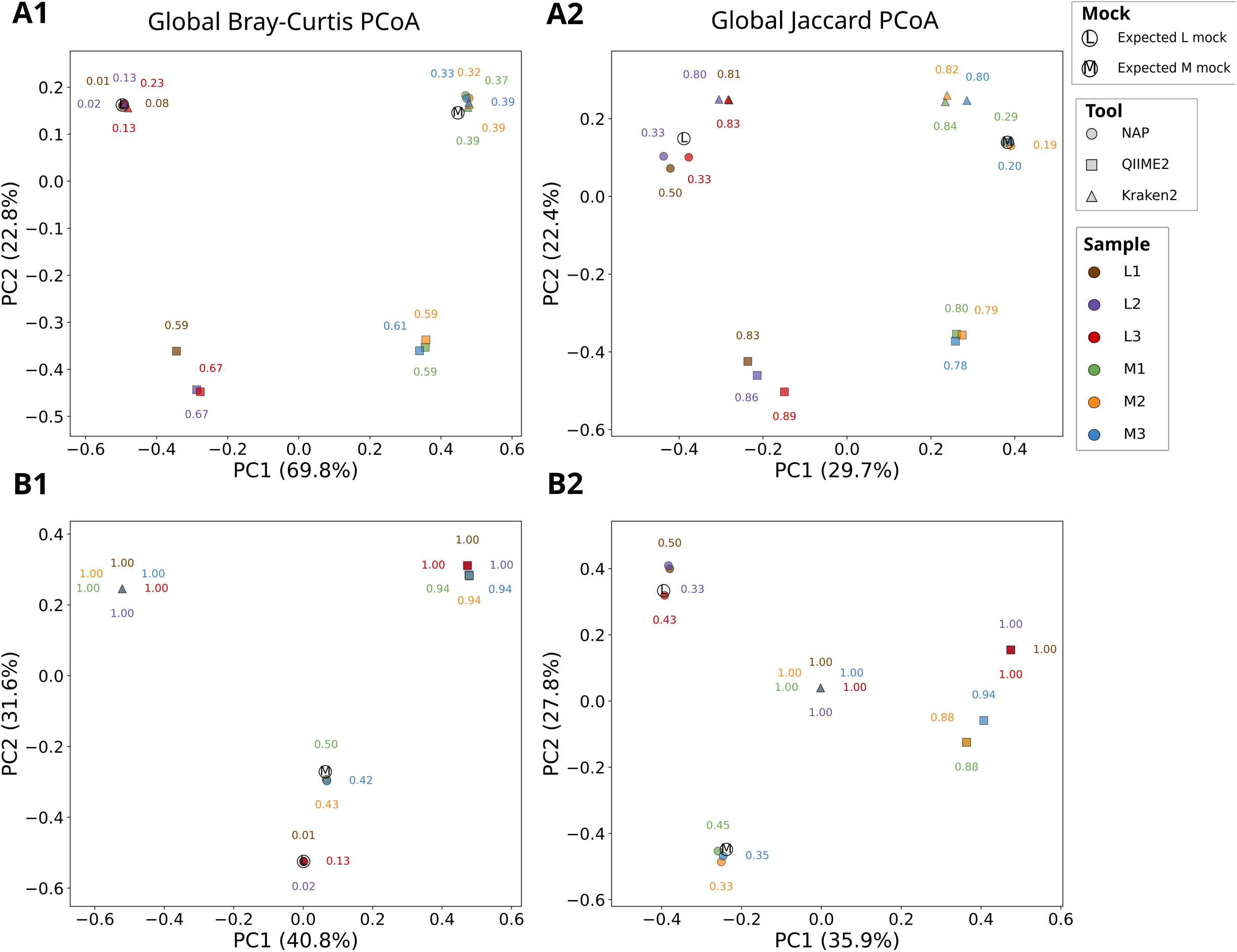
Principal coordinate analysis of samples *per* benchmarking tool. (A) Genus level results. (B) Species level results. Rows (1) and (2) shows Bray-Curtis and Jaccard metrics accordingly.

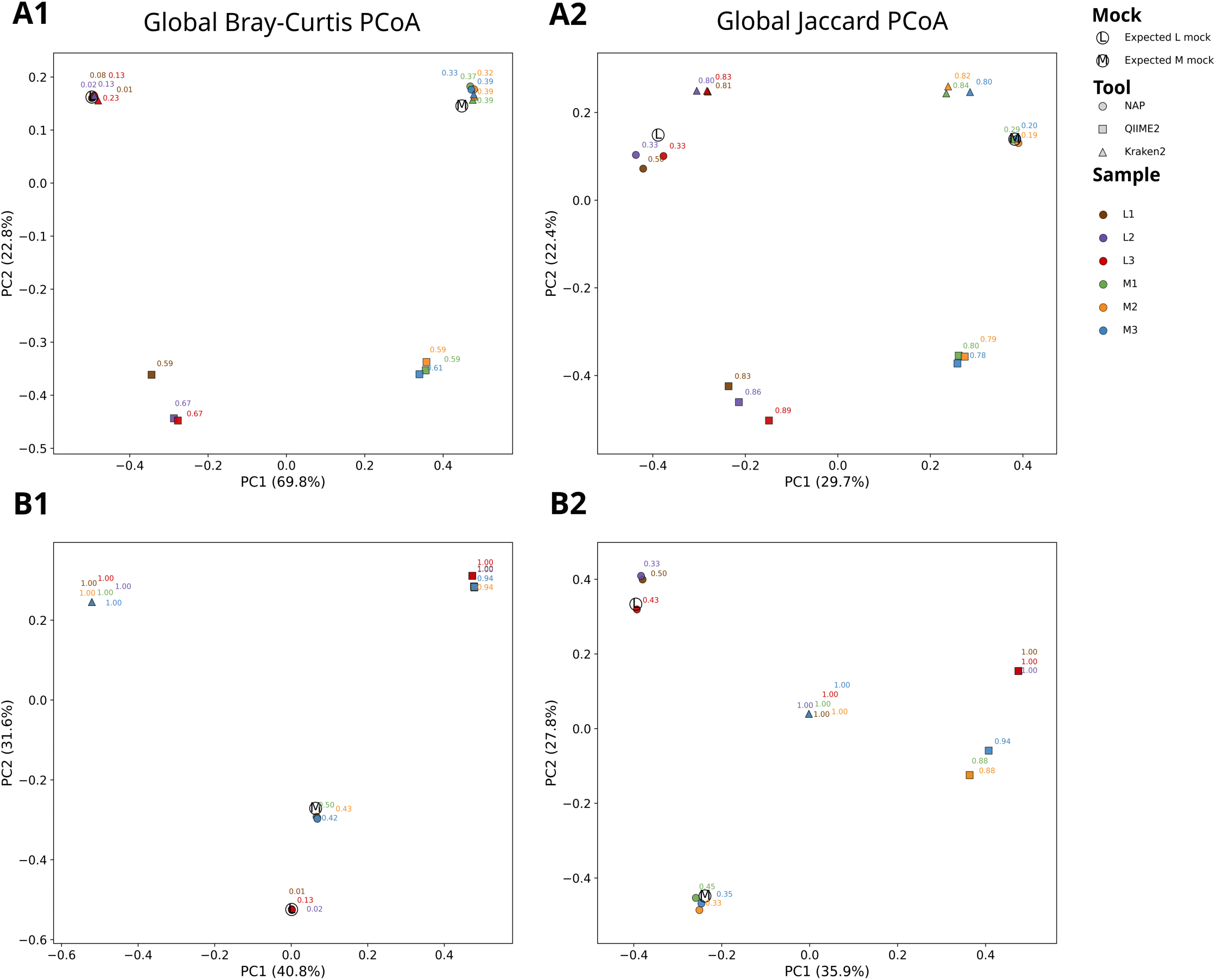

### NAP-based taxonomic classification

All expected bacterial and eukaryotic genera were detected across both mock communities, with only *Saccharomyces* exhibiting high replicate variability (coefficient of variation > 0.5; Figure 4A). The only archaeal genus (*Methanobrevibacter*) was detected below the pipeline’s 0.05% relative abundance filtering threshold. Similarly, *Salmonella* was detected but also fell below the applied filtering threshold. Only four genera were identified as false positives at the genus level: *Shigella*, likely reflecting misclassification due to the presence of multiple *Escherichia* strains and their known sequence similarity within the V4–V5 small subunit ribosomal RNA regions; and *Bysmatrum*, *Pasteurellaceae*, and *Streptococcus*, which displayed sporadic, highly variable abundances and were also present in blank controls, suggesting contamination beyond the detection limits of the current decontamination method. This interpretation is supported by previous reports identifying *Lactobacillus, Saccharomyces, and Streptococcus* as common laboratory- or human-associated contaminants (Glassing *et al*., 2016; Salter *et al*., 2014).

Taxonomic accuracy was mostly preserved at the species level. However, *Veillonella rogosae* was not detected in any gut mock replicates; instead, artefactual misattribution led to identification of *V. dispar* and *V. parvus*. A similar but less pronounced pattern was observed for *Lactobacillus fermentum*, which showed inflated abundance as *L. acidophilus*, a known and present contaminant, was identified as *Lactobacillus fermentum*. These cases are consistent with the high sequence similarity of the V4–V5 region among closely related taxa (Janda and Abbott, 2007). For both *Veillonella* and *Lactobacillus*, minor Nanopore sequencing errors likely contributed to taxonomic divergence during species level classification.

At both genus and species levels, replicate profiles showed statistically significant fidelity to the expected mock community composition (*p* < 0.05; Figure 4), indicating high reproducibility and classification accuracy. Most expected taxa were successfully identified with low inter-replicate variability (e.g., CV < 0.4 in most cases, often reaching < 0.15). Contaminants were identified based on prevalence in blank controls, high blank-to-sample abundance ratios, and exclusion from expected taxa lists. On average, the decontamination process removed 7.00 ± 2.68 species level hits *per* replicate, all of which were present in blank samples and judged to be contaminants following manual review. An additional 9.83 ± 6.49 species *per* replicate exhibited altered abundances after decontamination without being fully removed. For several high abundance taxa (such as *Faecalibacterium prausnitzii, Listeria monocytogenes*, and *Pseudomonas aeruginosa),* these adjustments moved observed abundances closer to expected values and reduced variability across replicates (Figure 4). Together, these findings indicate that, although the current decontamination strategy is relatively simple and does not fully resolve artefactual misclassification, it effectively identifies common contaminants and improves taxonomic accuracy. Classification performance is highly reliable at the genus level, and with improved primer design or longer read lengths, species level accuracy could be enhanced.

### NAP-based taxon abundance accuracy

The pipeline’s default primers, 515Y/926R, target the V4–V5 region of the small subunit (SSU) rRNA and offer broad cross-domain coverage, amplifying *ca.* 96% of known rRNA sequences across bacteria, archaea, and eukaryotes. Although these primers improve upon earlier cross-domain designs, they still introduce bias through differential binding site specificity and annealing efficiency, which can skew taxon representation (Parada *et al*., 2016; McNichol *et al*., 2021). Such effects are further compounded by wet lab variables, particularly DNA extraction efficiency, and SSU copy number variation. To mitigate these biases, we implemented prolonged bead beating cycles (Detailed in ‘Mock community preparation’) to balance DNA recovery from both easily lysed Gram-negative bacteria and more resilient taxa, including Gram-positive bacteria, spore formers, and eukaryotes (Zhang *et al*., 2020).

We assessed the accuracy of relative abundance estimates by comparing observed taxon abundances against expected values from mock community profiles using agreement plots and Bland–Altman analysis at genus level (Figure 6). The logarithmic mock community showed excellent concordance, with minimal Bland–Altman bias (21.2 units) and strong statistical agreement across replicates (*r* = 1, *p* = 1.5e^−5^, *p*_c_ = 1). In contrast, the gut mock community showed lower correlation (*r* = 0.68, *p* = 2.5e^−3^, *p_c_* = 0.53) and a substantial negative bias (–5876.6), suggesting systematic underrepresentation of several expected taxa. However, intra-taxon variability remained low (Figure 4A), indicating that discrepancies were driven by consistent biases, potentially due to primer mismatch or extraction inefficiencies, rather than random noise or computational error. To further validate the pipeline’s reliability, we repeated the agreement and Bland–Altman analyses at species level (Figure 7). Results remained statistically robust, with only marginal reductions in correlation (e.g., gut mock: *r* = 0.68 genus vs. 0.66 species; *p* = 2.5×10⁻³ vs. 1.7×10⁻³; *ρ_CCC_* = 0.53 vs. 0.56) and reduced Bland–Altman bias in both mock communities (e.g., log mock: 21.2 genus vs. 17.7 species). These patterns suggest that, in well characterised taxa, species level profiles produced by the pipeline remain quantitatively trustworthy. Nonetheless, while species level abundances appear numerically more concordant, taxonomic resolution is often compromised by limitations of the V4–V5 region, which can fragment expected taxa into multiple spurious species (e.g., *Veillonella* being misassigned to two absent species). As such, genus level assignments remain more biologically reliable than species level, and are therefore preferred for robust interpretation under the pipeline’s default settings.

**Figure 6:**
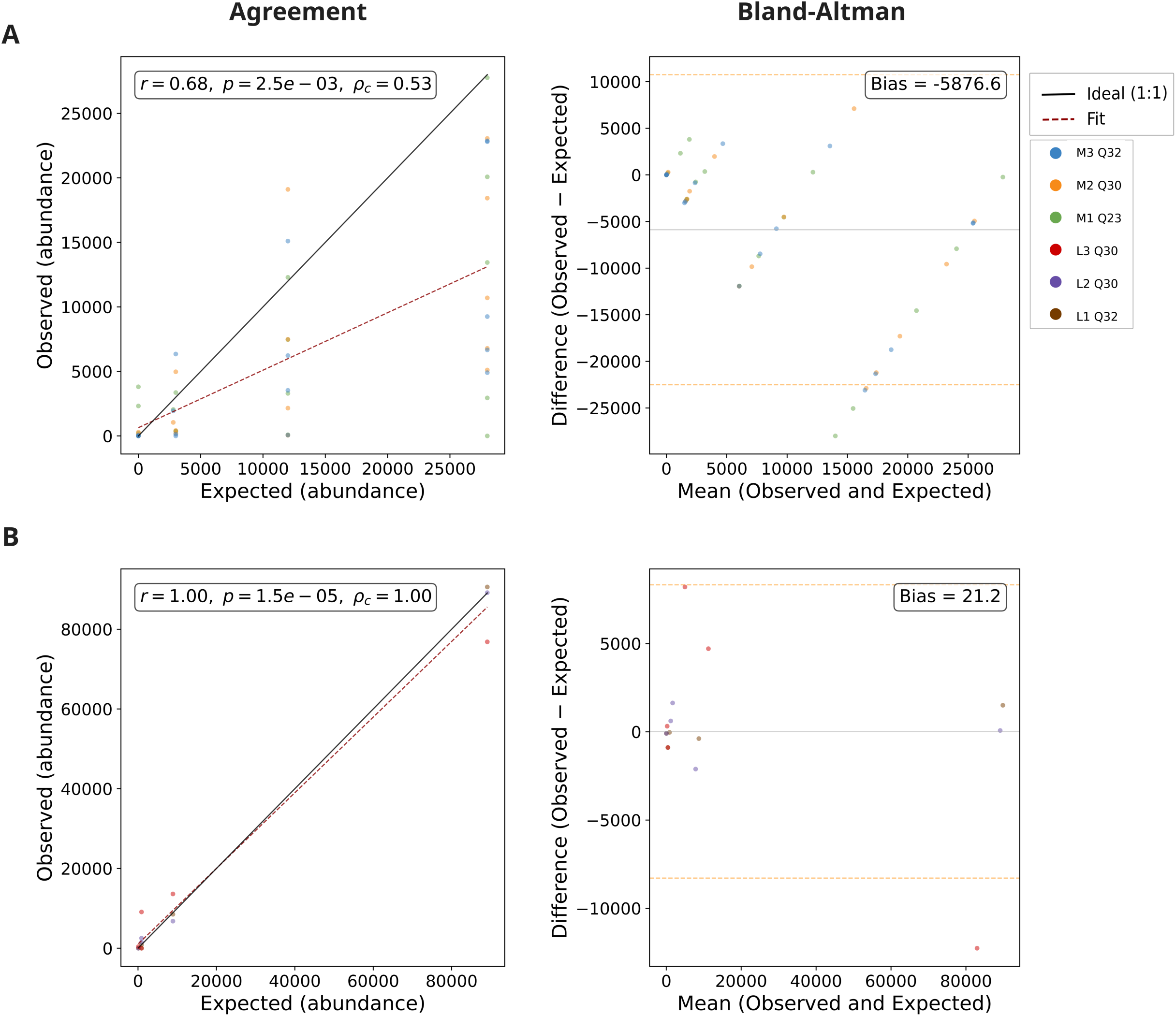
Agreement and error in taxonomic quantification across mock communities at genus level. (A) Gut mock community (M1–M3). (B) Logarithmic mock community (L1–L3). Left-hand plots show agreement between expected and observed *per* taxon genus level abundances. Agreement plots are annotated with Pearson correlation coefficient (*r*), p-value (*p*), and Lin’s concordance correlation coefficient (*ρ*_c_). Right-hand plots show Bland–Altman analysis, revealing mean bias and spread of differences across replicates, with bias annotated. This analysis (both agreement and Bland-Altman) only includes taxa present in the expected profile and their corresponding observed abundances; false positives are excluded.

**Figure 7:**
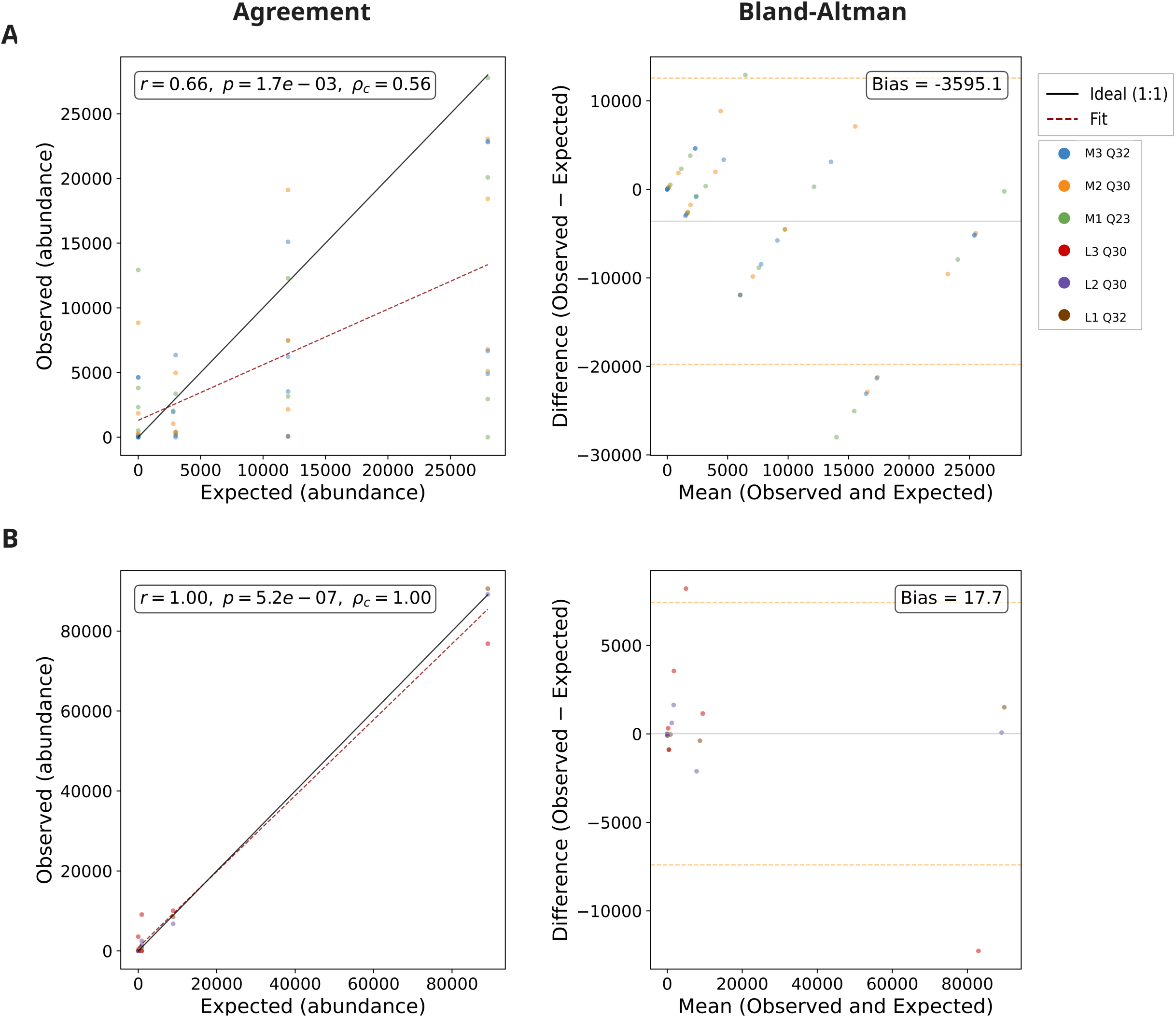
Agreement and error in taxonomic quantification across mock communities at species level. (A) Gut mock community (M1–M3). (B) Logarithmic mock community (L1–L3). Left-hand plots show agreement between expected and observed *per*-taxon species level abundances. Agreement plots are annotated with Pearson correlation coefficient (*r*), p-value (*p*), and Lin’s concordance correlation coefficient (*ρ*_c_). Right-hand plots show Bland–Altman analysis, revealing mean bias and spread of differences across replicates, with bias annotated. This analysis (both agreement and Bland-Altman) only includes taxa present in the expected profile and their corresponding observed abundances; false positives are excluded.

While 515Y/926R taxonomic bias has not been well explored, there is evidence that the observed abundance discrepancies are more consistent with combined wet lab and marker-specific bias than with classification failure alone. The phylum *Pseudomonadota* (formerly *Proteobacteria*) was well represented, with *Pseudomonas* showing expected abundances, as observed in other studies (Klindworth *et al*., 2013). Conversely, as expected, *Actinobacteriota* was underrepresented, with *Bifidobacterium* displaying particularly low abundance. Amplicon bias is known in this phylum, and is potentially compounded here by the Gram-positive cell wall structure (Parada *et al*., 2016). Bacillota (previously Firmicutes) were moderately underrepresented overall, while Bacteroidetes were more substantially affected, a pattern of particular interest given the clinical relevance of the ‘Firmicutes:Bacteroidetes’ ratio (Palkova *et al*., 2021). Within these phyla, *Faecalibacterium*, *Lactobacillus, Listeria*, and *Veillonella* showed good or mildly reduced recovery, while *Bacteroides* and *Prevotella* were more strongly underrepresented. This skew was less pronounced than in some prior studies, possibly due to differential lysis efficiency: Gram-positive Bacillota may be more effectively extracted than Gram-negative Bacteroidetes under prolonged bead beating conditions (Parada *et al*., 2016; Zhao *et al*., 2023).

*Bacillus* detection was inconsistent across replicates. In two gut mock community samples, it was present below the reporting threshold; in a third, it appeared overrepresented. As a spore forming, Gram-positive taxon, *Bacillus* is known to be sensitive to extraction and amplification biases. Additionally, Table 1 supports the possibility of mismatch-based primer bias, a key factor in under-amplification of rRNA. Furthermore, its presence in extraction blanks suggests potential cross-contamination, meaning its abundances are further reduced. Misalignment may also explain the inflated abundance in one replicate, given the high sequence similarity between *Bacillus* strains, where a single significant contamination event may have reinforced the taxon’s apparent presence. A comparable pattern was seen with *Shigella*, which was not present in the mock community but appeared alongside *Escherichia*, likely due to shared rRNA sequence regions and minor contamination (Zhao *et al*., 2023).

**Table 1:** *In silico* TestPrime summary of 515Y/926R primer compatibility for taxa represented in the mock communities. Analysis used SILVA SSU RefNR 138.2 with up to three mismatches and no enforced zero mismatch zone at the 3′ end. “Perfect Match” indicates the percentage of unambiguous primer–reference matches with zero mismatches; “Mean Pair Coverage” indicates overall primer-to-template coverage across best unambiguous matches; “3′-end MM Forward” and “3′-end MM Reverse” indicate the percentage of matches containing at least one 3′-end mismatch in the respective primer.

| Species | Perfect Match (%) | Mean Pair Coverage (%) | 3'-end MM Forward (%) | 3'-end MM Reverse (%) |
| --- | --- | --- | --- | --- |
| *Akkermansia muciniphila* | 100.0 | 100.0 | 0.0 | 0.0 |
| *Bacillus subtilis* | 94.2 | 99.7 | 2.4 | 1.9 |
| *Bacteroides fragilis* | 100.0 | 100.0 | 0.0 | 0.0 |
| *Bifidobacterium adolescentis* | 100.0 | 100.0 | 0.0 | 0.0 |
| *Candida albicans* | 92.9 | 99.6 | 2.9 | 0.0 |
| *Clostridioides difficile* | 99.6 | 100.0 | 0.4 | 0.0 |
| *Escherichia coli* | 99.7 | 100.0 | 0.1 | 0.1 |
| *Faecalibacterium prausnitzii* | 100.0 | 100.0 | 0.0 | 0.0 |
| *Fusobacterium nucleatum* | 97.8 | 99.9 | 0.7 | 0.7 |
| *Lactobacillus fermentum* | 95.4 | 99.8 | 3.4 | 0.6 |
| *Listeria monocytogenes* | 99.3 | 100.0 | 0.4 | 0.1 |
| *Methanobrevibacter smithii* | 100.0 | 100.0 | 0.0 | 0.0 |
| *Prevotella corporis* | 100.0 | 100.0 | 0.0 | 0.0 |
| *Pseudomonas aeruginosa* | 95.7 | 99.8 | 1.7 | 1.5 |
| *Roseburia hominis* | 100.0 | 100.0 | 0.0 | 0.0 |
| *Saccharomyces cerevisiae* | 91.4 | 99.3 | 2.9 | 0.0 |
| *Salmonella enterica* | 99.7 | 100.0 | 0.1 | 0.1 |
| *Veillonella rogosae* | 100.0 | 100.0 | 0.0 | 0.0 |

Other taxa have poorly characterised biases for primer set 515Y/926R, limiting interpretation. Here, for example, *Methanobrevibacter smithii* and *Salmonella enterica* were both detected below the pipeline’s filtering threshold (0.05% relative abundance filtration threshold; 0.2% and 0.089% expected relative abundance for *M. smithii* and *S. enterica*, respectively), suggesting mild underrepresentation, with only *Salmonella enterica* showing potential for primer mismatches (Table 1). Similarly*, Akkermansia muciniphila* showed no primer mismatches and underrepresentation, with bias uncharacterised in the literature. For *Methanobrevibacter smithii*, the difficulty of archaeon lysis is a strong candidate contributor, but other factors can lead to related archaea being underrepresented (Youngblut *et al.,* 2021; Zhao *et al*., 2023). *Akkermansia muciniphila* and *Salmonella enterica* are easy to lyse Gram-negative, non-spore forming bacteria, suggesting excessive bead beating may have led to a slight underrepresentation bias, but this is likely compounded by other PCR-related biases, considering mismatches are not the only known factor in primer-induced bias (Qin *et al*., 2023; Silverman *et al*., 2021).

Finally, as the 515Y/926R primers also capture eukaryotic 18S rRNA, a domain correction factor is required to adjust for lower 18S amplification efficiency. The pipeline applies a default correction factor of 0.4, based on empirical estimates ranging from 0.3–0.5 (Yeh *et al*., 2021). This correction was validated in our results: *Candida albicans* and *Saccharomyces cerevisiae* were detected at or near to the expected abundances relative to co-occurring 16S taxa.

Taken together, replicate abundances were highly consistent, demonstrating strong technical reproducibility. Deviations from expected values were largely attributable to primer mismatch, variability in lysis efficiency, or low input abundance (rather than stochastic or computational error). The pipeline’s default 16S/18S correction proved effective, and the bead beating protocol did not seem to induce a systematic skew between Gram-positive and Gram-negative bacterial taxa. These findings support the pipeline’s capacity to generate biologically representative taxonomic profiles under the tested conditions.

### Quantitative performance metrics

Detection sensitivity and classification accuracy were assessed across the logarithmic and gut mock communities at both genus and species levels (Figure 8). For NAP, expected taxa above *ca.* 1% relative abundance were generally detected consistently across replicates, whereas recovery below this level became progressively less stable. At genus level, all expected taxa at 8.9% and 89.1% in the logarithmic mock were recovered in all replicates, while the lower abundance taxa at 0.89% and 0.089% showed variable or absent recovery. In the gut mock, most expected taxa at ≥1.5% were detected in all three replicates, with the main exceptions being *Candida* and *Roseburia* (both 2/3 replicates) and *Methanobrevibacter*, which was not recovered above filtering threshold (0.05%). Species-level detection showed the same overall trend, although recovery of some closely related taxa became less stable, most notably *Veillonella rogosae*, which was not recovered above threshold despite an expected abundance of 14%.

**Figure 8:**
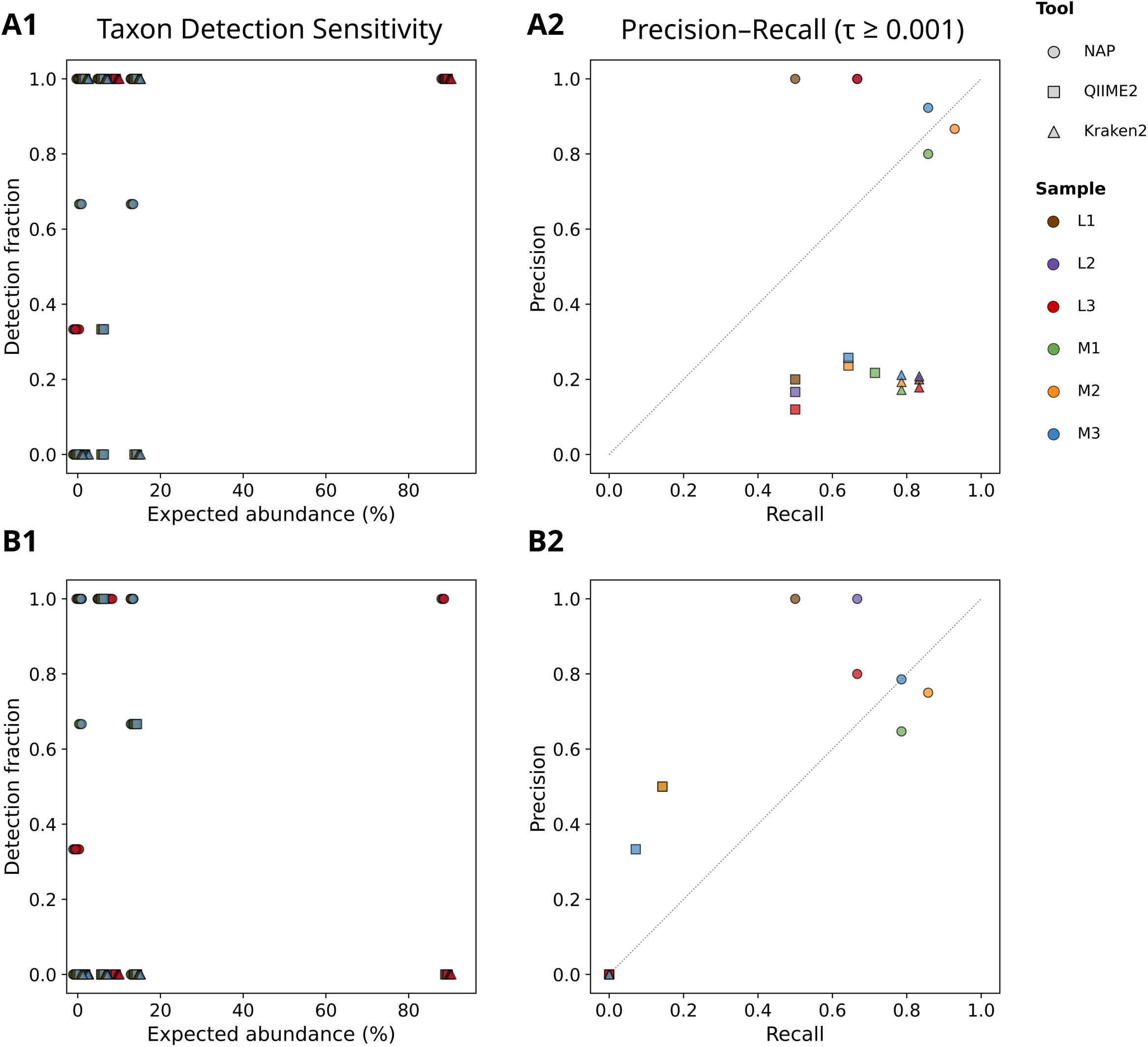
Detection sensitivity and precision–recall performance across benchmarked workflows. (A) Genus-level and (B) species-level resolution shown, with logarithmic and gut mock community results combined. Left: detection sensitivity across expected taxa, where each point represents a taxon plotted by its expected relative abundance (%) and the fraction of replicates in which it was detected. Right: *per*-replicate precision–recall values calculated at threshold τ ≥ 0.001. Point shape indicates workflow and colour indicates sample identity.

Precision–recall analysis showed that NAP retained strong classification performance across both taxonomic ranks. At genus level, precision ranged from 0.80 to 1.00 and recall from 0.50 to 0.93 across the six replicates, corresponding to mean precision, recall, and F1-score values of 0.93, 0.75, and 0.81, respectively. Species-level performance declined only modestly, with precision ranging from 0.65 to 1.00 and recall from 0.50 to 0.86, yielding mean values of 0.83, 0.71, and 0.75. Relative to the benchmarked alternatives, NAP also retained substantially stronger precision at genus level and was the only workflow to preserve meaningful species-level precision–recall behaviour under the applied threshold. QIIME2 showed reduced genus-level precision due to numerous unexpected taxa and markedly weaker species-level performance (mean precision 0.22, recall 0.06, F1-score 0.19), whilst Kraken2 did not recover species-level true positives above threshold.

Taken together, these results indicate that NAP provides strong classification accuracy and consistent detection sensitivity across replicate samples, particularly at genus level. Recovery was most reliable above *ca.* 1% relative abundance, with increasingly stochastic detection below this level, especially in the logarithmic mock. Under the applied threshold, the lowest abundance taxon recovered in the logarithmic mock was detected at 0.089% in one of three replicates, whereas the lowest abundance taxon recovered consistently in the gut mock was present at 1.5%. These data therefore place the practical detection range for the present assay between *ca.* 0.1% and 1% relative abundance, although it cannot be defined more precisely from the available mock communities. In the context of the direct benchmark, NAP also maintained substantially stronger genus-level precision than QIIME2 and Kraken2 using the same data, and remained the only workflow to preserve meaningful species-level sensitivity and F1 performance, together with precision and recall consistently in line with established performance expectations for amplicon-based microbiome profiling (Poncheewin *et al*., 2020). Moreover, the observed precision and recall values compare favourably with Illumina-based profiling, using workflows such as NG-Tax and QIIME2 (Poncheewin *et al*., 2020), highlighting the pipeline’s reliability. These findings support the utility of NAP for accurate, reproducible, and domain-inclusive microbiome profiling under the tested conditions.

## Discussion

Under the conditions tested here, NAP recovered the expected structure of two complementary mock communities with good reproducibility and strong genus-level fidelity, while also comparing favourably with QIIME2 and Kraken2/Bracken on the same input data. The clearest differences emerged in false positive behaviour and in the stability of species-level outputs. At genus level, the clearest advantage of NAP was the marked reduction in unexpected assignments relative to QIIME2 and Kraken2/Bracken. This behaviour is consistent with the centroid-first and consensus-based design of NAP, which likely limits propagation of Nanopore read errors into downstream taxonomic classification by reducing reliance on noisy single-read assignments. At species level, this difference became more pronounced, with NAP retaining informative output where the comparator workflows deteriorated sharply under the present conditions. Taken together, these results support NAP as a practical workflow for cross-domain Nanopore amplicon profiling, with its strongest performance currently at genus level.

The two mock communities used here were chosen to test different aspects of performance rather than to represent the full diversity of natural microbiomes. The logarithmic mock provided an extreme abundance gradient and therefore challenged abundance-dependent detection, whereas the gut mock imposed greater compositional complexity and included taxa expected to vary in extractability, amplification behaviour, and separability within the V4–V5 region. Together they provided bacterial, archaeal, and eukaryotic targets and allowed assessment of cross-domain recovery, abundance fidelity, and reproducibility under controlled conditions. They do not, however, capture the full taxonomic and compositional diversity of environmental or host-associated samples, and the present validation should therefore be interpreted as establishing core performance rather than universal applicability across all microbiome types.

Several features of NAP likely contributed to the behaviour observed here, including reduced taxonomic artefacts and improved contamination handling (*via* reduction in false positives), greater stability across variable read inputs, and more accurate preservation of expected community structure. The workflow combines primer-aware database refinement, centroid-based sequence compression, BLAST-based taxonomic assignment, hierarchical consensus correction, and RAW-read reassignment to an internal high confidence reference set. In practice, this appears to reduce the impact of noisy single read classifications while preserving abundance information and limiting false positive inflation. This interpretation is consistent with the direct benchmark: the principal advantage of NAP was not simply higher recovery of expected taxa, but cleaner preservation of expected community structure once unexpected detections were taken into account. In that sense, the workflow’s contribution is primarily one of integration and control of error propagation across an end-to-end analysis path for a data type (Nanopore sequencing) that currently lacks a well established cross-domain solution.

NAP also demonstrated robust performance across variable sequencing conditions. The adaptive filtering benchmark showed that the default filtering configuration was not the single numerical optimum for every summary metric, but it was the most reliable compromise across mixed read depths and qualities. Fixed Q30 filtering was clearly too stringent, particularly for low input samples, whereas aggressive masking degraded downstream taxonomic fidelity. Fixed Q20 modes improved some sensitivity-oriented endpoints, but only by applying the same permissive threshold to all samples and abandoning sample-specific adaptation. The default dynamic strategy therefore appears to function as intended, preserving sufficient depth for lower input samples while maintaining stricter filtering where depth permits. This is reflected in the classification metrics: across the six NAP replicates, mean genus-level precision, recall, and F1-score were 0.93, 0.75, and 0.81, respectively, declining only modestly at species level to 0.83, 0.71, and 0.75. These metrics were accompanied by consistently high read classification rates, typically exceeding 99.6%, with only modest reductions observed in lower depth datasets. Overall, this level of performance is comparable to that reported for established Illumina-based amplicon pipelines, such as NG-Tax (Ramiro-Garcia *et al*., 2018). Detection was most reliable above approximately 1% relative abundance, with increasingly unstable recovery below this level; with the present assay, the practical detection range therefore appears to lie somewhere between *ca*. 0.1% and 1% relative abundance. These values are comparable to those reported for established Illumina-based amplicon workflows, which similarly balance precision and recall in mock community benchmarking (Poncheewin *et al*., 2020). Consistent with this, genus-level Bray–Curtis dissimilarities were low in the logarithmic mock community (0.01–0.13), within the range of technical variation reported for mock community sequencing (Yeh *et al*., 2017), and remained moderate in the more complex gut mock community (0.32–0.39). Overall, this level of variation is within the 0.2–0.5 range observed across state-of-the-art pipelines in inter-laboratory benchmarking studies, where differences in analytical approach alone can influence both taxon detection and relative abundance estimates (as a proxy for the variance expected within a single pipeline’s outputs; O’Sullivan *et al*., 2021).

A second important finding is that the remaining disagreement between observed and expected abundances appears to arise largely from assay-level limitations rather than computational instability. This was most evident in the gut mock, where several expected taxa were consistently underrepresented despite relatively low within-group variability. Such behaviour is more consistent with systematic bias than with erratic classification. The taxa affected, and the directions of bias observed, also fit with known issues in amplicon profiling, including primer-template mismatch, differences in lysis efficiency, contamination, and limited taxonomic separability within short ribosomal regions (Parada *et al*., 2016; Janda and Abbott, 2007; Silverman *et al*., 2021). The *Escherichia–Shigella* and *Veillonella* examples illustrate this clearly: abundance may still be recovered in approximately the right part of taxonomic space, but fine-scale assignment can fragment across closely related alternatives. The fact that broad abundance distortions were also visible in the comparator workflows supports the view that these discrepancies are not unique to NAP, but reflect constraints of the assay and marker region itself.

The same reasoning helps explain why genus-level interpretation is currently the strongest use case for the workflow. Across the present data, genus-level profiles were reproducible and closely matched expected community structure, whereas species-level results were more conditional. Some taxa remained well resolved at species level, but others did not, even where abundance concordance appeared numerically acceptable. For the default 515Y/926R amplicon, species-level outputs should therefore be treated as informative where supported, rather than assumed to be uniformly reliable. The decontamination step improved several profiles and reduced the impact of likely contaminants, but it did not fully resolve all artefactual assignments, particularly where contamination and limited marker resolution interacted.

There are several further limitations that should be made explicit. First, validation was limited to two mock community systems and did not include field-derived or clinical samples. Second, the practical limit of detection in this assay appears to lie somewhere between approximately 0.1% and 1% relative abundance, with reliable recovery becoming much less stable below the upper end of that range. Third, NAP does not currently apply taxon-specific SSU copy number correction. In a cross-domain context, robust copy number adjustment remains difficult because curated information is incomplete and uneven across bacteria, archaea, and eukaryotes. Output abundances should therefore be interpreted primarily as relative amplicon abundances rather than direct estimates of cellular abundance. Finally, although the workflow was benchmarked here against QIIME2 and Kraken2/Bracken, broader comparison across additional datasets and primer systems would be valuable in defining where the present approach functions well and where further tuning is required.

In conclusion, NAP provides an accurate, reproducible, and open source workflow for long-read, cross-domain microbiome profiling under the tested conditions. Its strongest performance is at genus level, where it combined good taxonomic fidelity with substantially lower false positive detection than the benchmarked alternatives, whilst maintaining mean precision, recall, and F1-score values of 0.93, 0.75, and 0.81 across the six replicates. Species-level outputs remained informative for some well resolved taxa, but were less robust overall and should be interpreted cautiously under the default V4–V5 assay. NAP does not remove the known constraints of amplicon profiling, including primer bias, extraction bias, copy number effects, and limited low abundance sensitivity, but it does provide a practical end-to-end workflow that preserves expected community structure well and supports mixed domain analysis from a single Nanopore amplicon dataset. On that basis, NAP appears well suited to studies seeking Nanopore-sequencing-compatible, domain-inclusive ribosomal profiling, provided that abundances are interpreted as relative amplicon abundances and that species-level conclusions are drawn with appropriate caution.

## List of abbreviations

ASV: amplicon sequence variant
bp: base pair
BLAST: basic local alignment search tool
CD-HIT: cluster database at high identity with tolerance
CNT: Centroid dataset
CV: coefficient of variation
DNA: deoxyribonucleic acid
HAC: High accuracy consensus
PCoA: principal coordinate analysis
PCR: polymerase chain reaction
Q-score (Phred score): quality score representing probability of error in base calling
rRNA: ribosomal ribonucleic acid
SSU: small subunit (of rRNA)
TSS: total sum scaling
TSV: tab-separated values
ρc (ρCCC): Lin’s concordance correlation coefficient, β-diversity: beta diversity

## Declarations

### Availability of data and materials

The datasets and code supporting the conclusions of this article are available as follows:

Project name: NAP

Project home page: https://github.com/Luke-B-Jones/NAP

Archived version: A permanently archived release is available *via* Zenodo along with early stage development outputs at DOI:10.5281/zenodo.19671570

Operating system(s): Platform-independent (tested on Linux)

Programming language: Python (≥3.9) and Bash

Other requirements: Docker, or a Standard UNIX environment

License: MIT license (open-source; free for academic and commercial use)

Restrictions to use by non-academics: None.

The raw Nanopore amplicon sequencing data generated during this study are available in the NCBI Sequence Read Archive (SRA) under BioProject accession PRJNA1367045.

### Authors’ contributions

LJ designed the study and implemented the pipeline, and carried out analyses. SB acquired funding and supervised the project. LJ and SB contributed to manuscript preparation. Both authors read and approved the final manuscript.

## Acknowledgements

We thank Adrien Leger, Oxford Nanopore Technologies Ltd, for critical reading of the manuscript. We thank Josephine Ilott and Morgan Cockrill, Department of Life Sciences, University of Bath, for their contributions to the NAP pipeline: development of the decontamination module and improvement of the pipeline’s error reporting and overall robustness, respectively. Both contributions have been documented in the project repository.

