## Supplemental Info 1 for "NAP: an open-source pipeline for cross-domain microbiome profiling using Nanopore sequencing-derived amplicon data"

### Supplementary Materials

During development of NAP, multiple clustering, consensus, and classification strategies were evaluated to identify a final workflow that preserved expected community structure whilst remaining computationally practical for Nanopore amplicon data. Candidate clustering approaches included CD-HIT, isONclust, and Rattle. Consensus-generation strategies included alignment-based polishing and centroid-guided approaches, and taxonomic assignment was explored using BLAST-based and alternative classification frameworks. These development-stage comparisons were performed on the same mock-community datasets used in the main manuscript and were assessed primarily using β-diversity proximity to expected mock composition, taxonomic classification rate, and overall output consistency. These optimisation analyses supported the final workflow centred on CD-HIT clustering, BLAST-based taxonomic assignment, and hierarchical consensus correction, which showed the closest convergence to expected mock-community composition across the configurations tested (Supplementary Figure 1).

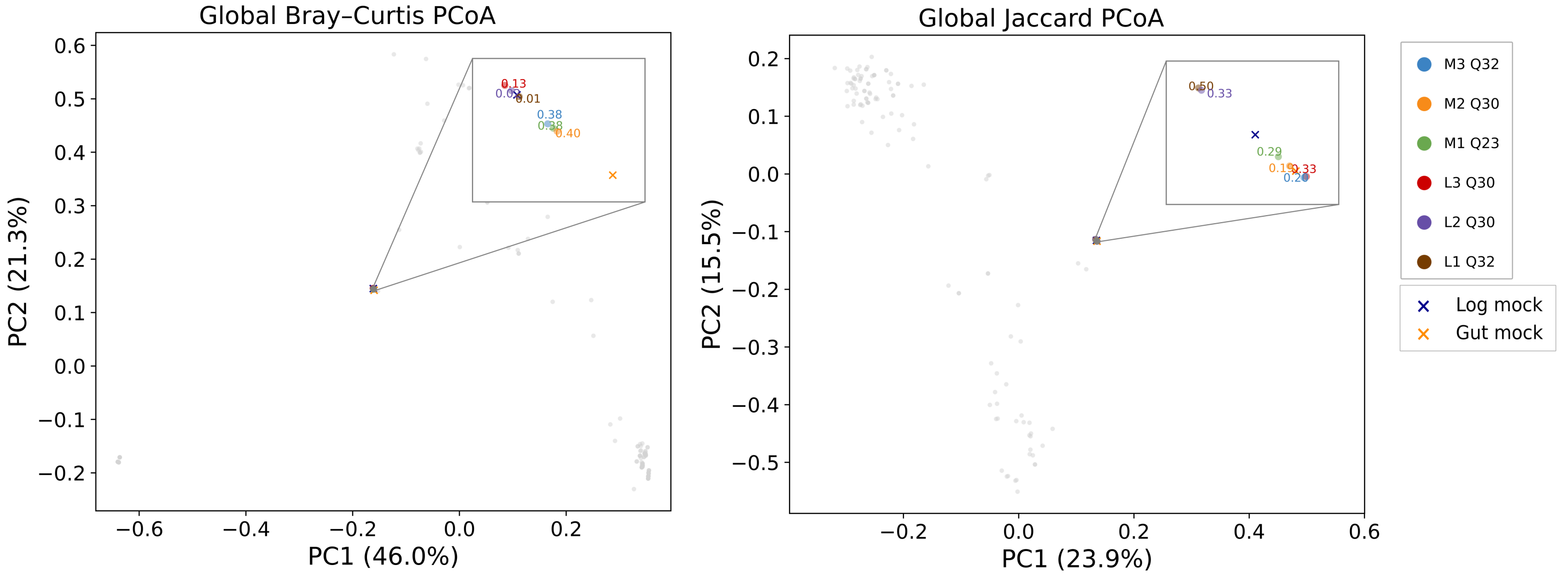


**Supplementary Figure 1:** Development-stage optimisation of the final NAP workflow. Grey points represent alternative clustering, consensus, and classification configurations evaluated during pipeline optimisation, plotted relative to the expected mock-community composition. Coloured points indicate the outputs of the final selected NAP configuration, and crosses indicate the expected mock profiles. Insets show the final selected outputs only, annotated with the corresponding β-diversity metrics. L1–L3 are replicates of the logarithmic mock community and M1–M3 are replicates of the gut mock community.

**Supplementary Table 1:** Default adaptive filtering thresholds used by the NAP configuration in this study. Samples were assigned to an input-read-count bin, and each bin was linked to a preset mean read Phred threshold and unmuted fraction target. The unmuted fraction defines the proportion of highest-quality bases retained unmasked during base-level quality masking; bases falling below the derived masking cutoff were converted to `N`. This adaptive scheme allows more stringent filtering in higher-depth datasets while preserving usable sequence in lower-depth samples

| Bin (input reads number) | Phred applied | Unmuted fraction |
|---|---|---|
| >2,000,000 | Q35 | 99.9% |
| >1,500,000 | Q33 | 99.9% |
| >500,000 | Q30 | 99.9% |
| >300,000 | Q28 | 99.8% |
| >200,000 | Q26 | 99.5% |
| >100,000 | Q25 | 99.5% |
| >50,000 | Q23 | 99.0% |
| ≤50,000 | Q20 | 99.0% |
